# tACS entrains neural activity while somatosensory input is blocked

**DOI:** 10.1101/691022

**Authors:** Pedro G. Vieira, Matthew R. Krause, Christopher C. Pack

**Affiliations:** Montreal Neurological Institute, McGill University, Montreal, QC H3A 2B4, Canada

## Abstract

Transcranial alternating current stimulation (tACS) modulates brain activity by passing electrical current through electrodes that are attached to the scalp. Because it is safe and non-invasive, it holds great promise as a tool for basic research and clinical treatment. However, little is known about how tACS ultimately influences neural activity. One hypothesis is that tACS affects neural responses *directly*, by producing electrical fields that interact with the brain’s endogenous electrical activity. Since the shape and location of these electric fields can be controlled, stimulation could be targeted at brain regions associated with particular behaviors or symptoms. However, an alternative hypothesis is that tACS affects neural activity *indirectly*, via peripheral sensory afferents. In particular, it has often been hypothesized that tACS acts on nerve fibers in the skin, which in turn provide rhythmic input to central neurons. In this case, there would be little possibility of targeted brain stimulation, as the regions modulated by tACS would depend entirely on the somatosensory pathways originating in the skin around the stimulating electrodes. Here, we directly test these competing hypotheses by recording single-unit activity in the hippocampus and visual cortex of monkeys receiving tACS. We find that tACS entrains neuronal activity in both regions, so that cells fire synchronously with the stimulation. Blocking somatosensory input with a topical anesthetic does not significantly alter these neural entrainment effects. These data are therefore consistent with the direct stimulation hypothesis and suggest that peripheral somatosensory stimulation is not required for tACS to entrain neurons.

## Introduction

Recent results suggest that transcranial alternating current stimulation (tACS) can non-invasively alter brain activity [1–4], but the physiological mechanisms behind these exciting findings remain poorly understood. Traditionally, tACS is thought to produce oscillating electric fields within the brain that hyperpolarize and depolarize neurons, so that they fire synchronously with the stimulation. Small animal experiments demonstrate that the fields generated by applying current to the bare skull can entrain neurons [1, 4, 5], consistent with intracranial electric fields having a *direct* effect of tACS on brain activity. In humans, however, the tACS electrodes are placed on the subject’s intact scalp, not within the skull. Since the skin is highly conductive, but the skull beneath is not, much of that current is shunted away from the brain and stimulates peripheral nerve fibers in the skin instead [6]. Rhythmic activation of these fibers could thus indirectly entrain central neurons by providing them with temporally structured sensory input. Since shunting also weakens electric fields in the brain, this *indirect* mechanism has been frequently proposed to be the dominant mode of action in humans [5, 7–10]. If this were true, it would have dramatic implications for how tACS is used and studied: brain areas would need to be targeted on the basis of somatosensory connectivity, rather than physical location, and brain regions that received little or no somatosensory input would be unreachable.

These competing hypotheses can be distinguished through the use of topical anesthesia. Pretreatment of the skin under and around the tACS electrodes with topical anesthetic blocks cutaneous afferents [11] and prevents them from generating somatosensory percepts [12]. If tACS acts indirectly via somatosensory inputs, topical anesthesia should reduce or abolish its effects by blocking transmission from the periphery. Conversely, if the electric fields directly affect neurons, applying topical anesthesia should produce little or no changes in the effects of tACS, as the same electric fields are produced within the brain in both conditions. Previous attempts to test the indirect hypothesis have used proxy measurements for neural activity, with mixed results: topical anesthesia appears to prevent tACS from affecting nociception [10] and tremor [5], but effects on motor-evoked potentials [13] and language processing [14, 15] persist when somatosensory inputs are blocked. Interpreting these results is challenging, because the neural mechanisms behind these readouts are not well-understood and each may involve multiple brain regions, only some of which may have been affected by the tACS used in each study.

An unambiguous test of the role of somatosensory input is to directly measure neural entrainment during tACS, with and without topical anesthetic. Here, we perform that decisive experiment in non-human primates, a highly realistic model for human neurostimulation. Using single-unit recordings of neurons in the hippocampus and visual cortex, we find that blocking somatosensory input has little effect on neural entrainment by tACS. Instead, our data support claims of a direct effect on neurons in the stimulated regions.

## Results

We collected data from two adult male rhesus monkeys (*Macaca mulatta*), using techniques and experiments that, with the exception of the topical anesthesia, are virtually identical to those in our previous work [3, 16]. Monkey N (7 year old male, 10 kg) participated in the experiments described in our previous study [3], while Monkey S (9 year old male, 20 kg) was obtained specifically for these experiments. These procedures were approved by the Montreal Neurological Institute’s Animal Care Committee and conformed to the guidelines of the Canadian Council on Animal Care.

First, we determined if 5% EMLA cream, a widely-used topical anesthetic, blocked the somatosensory stimulation produced by tACS. We used the animal’s behavioral performance to validate the effectiveness of this anesthesia. In our previous studies [3, 16], animals were initially distracted and agitated by the onset of tACS, but eventually learned to continue working despite the evoked percepts. If EMLA effectively blocks somatosensation, applying it around tACS electrodes should reduce these distractions and increase the time spent on task. Since a naive subject would be most sensitive to these effects, Monkey S, a well-trained monkey that had never received tACS, was tested using the paradigm in Fig 1A. Two pairs of tACS electrodes were placed on its head, one on anesthetized skin over one hemisphere and the other at identical locations on the untreated contralateral side. After a 20-minute delay, introduced to account for setup time needed in subsequent neurophysiology experiments, the monkey performed a visual fixation task while bursts of tACS were applied through each pair of electrodes.

**Fig 1.**
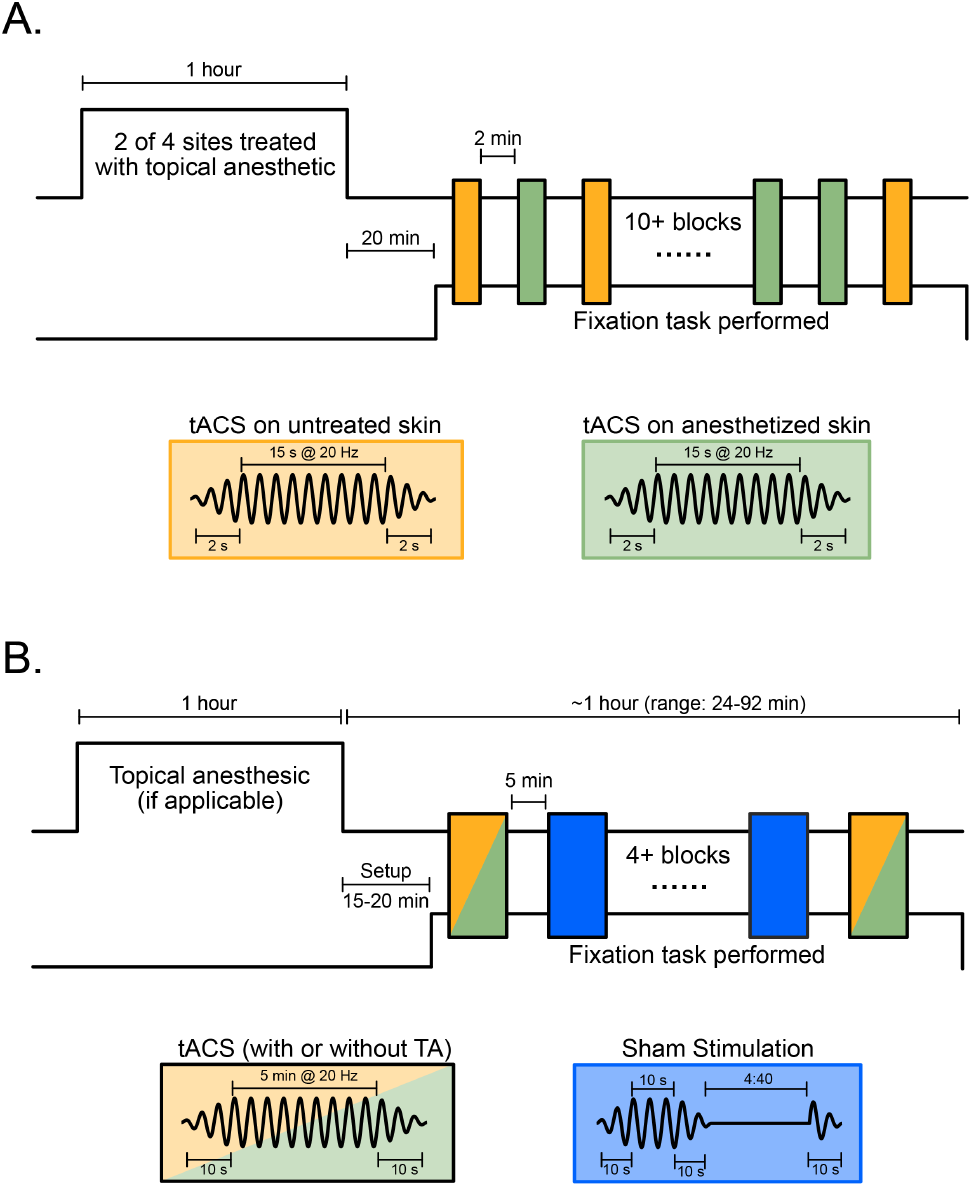
Experimental Design. **(A)** To measure the effectiveness of topical anesthesia, we applied short blocks of tACS through two pairs of electrodes, one of which was placed over skin pretreated with 5% EMLA cream (green); the skin under the other pair was left untreated (yellow). Since subjects typically adapt to tACS sensations, the short blocks and brief ramps maximize any behavioral effects of stimulation. **(B)** To record neural responses to tACS, longer blocks of stimulation were applied through a single set of electrodes placed at scalp locations that optimally simulated left hippocampus or left V4. In some sessions, the skin beneath both electrodes was pretreated with topical anesthetic (green); in others, the skin was left untreated as a control (yellow). In every session, a mixture of active tACS (yellow or green) and sham stimulation (blue), was applied in 5 minute blocks, with the ramp periods in increased to 10 seconds to reduce sensations. This design ensures that identical electric fields were generated during the topical anesthesia and control conditions. In both experiments, stimulation conditions were randomly interleaved and the tACS frequency was 20 Hz with amplitudes of ±0.5, ±1, or ±2 mA. Inset figures show the tACS waveforms in each condition.

As Fig 2 shows, time-on-task was significantly increased (p < 0.05) at all stimulation amplitudes when tACS was applied to the anesthetized skin, as compared to control sites. Furthermore, performance was not significantly different from ceiling (100%) during 0.5 and 1.0 mA tACS (0.5 mA: p = 0.07, t(10) = −2.01; 1 mA: p = 0.12; t(10) = −1.72; 1-sample *t*-test) with anesthesia, but was reduced during control conditions with somatosensation intact (p=0.01; t(10) = −3.13 and p < 0.001; t(10) −5.04, respectively). These data suggest that EMLA increases somatosensory thresholds to ~1 mA, and reduces—but does not eliminate—sensation at 2.0 mA, just as it does in humans [5].

**Fig 2.**
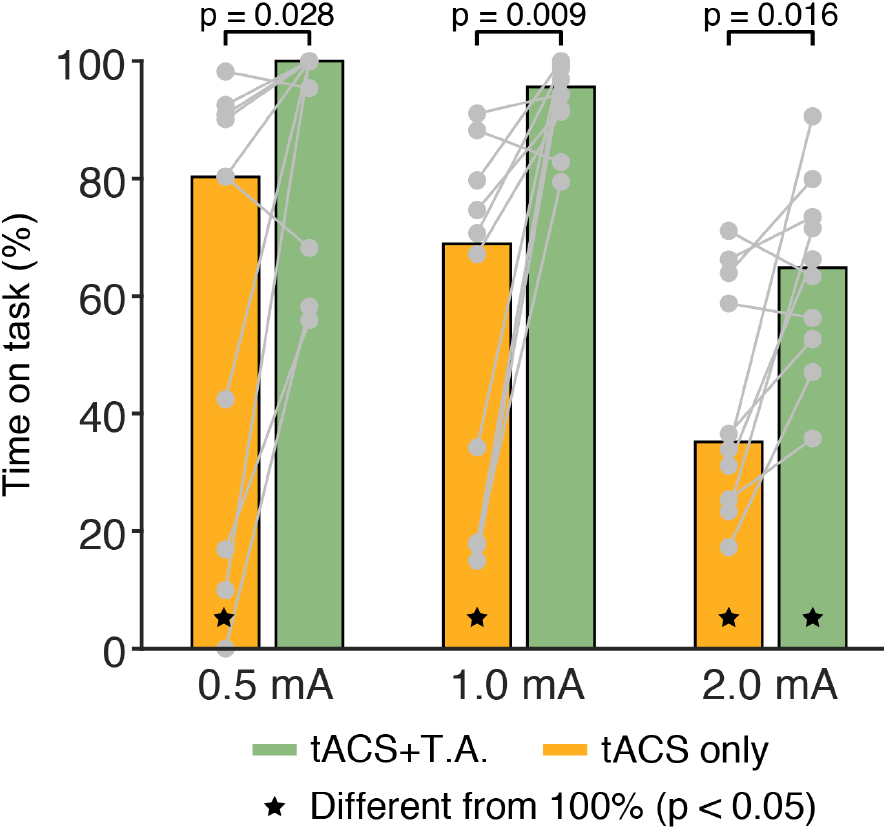
EMLA effectively blocks or reduces tACS-related somatosensation. In humans, tACS produces nociceptive sensations that disrupt behavior. To quantify those effects in our monkey subjects, the paradigm shown in Fig 1A was used. We recorded the proportion of time that the animal’s gaze remained within 2 degrees of the fixation target. Topical anesthesia (TA, green bars) significantly reduced behavioral disruptions at all current levels when compared to stimulation applied over intact skin (yellow bars). Individual data points are shown in grey; colored bars indicate the median. Black stars indicate conditions where performance was significantly below ceiling (100%; p<0.05)

Next, we recorded neural activity with and without somatosensory blockade. During some recording sessions, the skin under and around each tACS electrode was pre-treated with 5% EMLA; in others, the skin was left untreated as a control. This between-sessions design ensures that identical electric fields are produced during anesthesia and control conditions. In each session, interleaved blocks of 20 Hz tACS and sham stimulation (Fig 1B; *Methods*), were applied to the monkeys’ scalps. For each neuron, we calculated two phase-locking values (PLVs): one summarizing its entrainment to the tACS waveform and another quantifying its entrainment to the matching frequency component (20 Hz) of the local field potential during sham [3]. These data allow us to determine the proportion of neurons that become entrained by tACS during each anesthesia condition. Comparing the strength of entrainment (PLVs) across tACS conditions provides an additional measure of the effects of topical anesthesia.

We first obtained data from the hippocampal recording sites described in [3]. As in that study, the tACS electrode montage was optimized to stimulation the left hippocampus, so that a ±2 mA alternating current produced an electric field of ~0.3 V/m at the recording site. Under control conditions (Fig 3A, yellow circles, N = 8 sessions), tACS entrained 50 percent of the hippocampus neurons (N = 28/56; p < 0.05 per-cell permutation tests). The median PLV during tACS was 0.054 (95% CI of the median: [0.032 – 0.068]), significantly larger than that observed during sham blocks (median:0, CI: [0 – 0.006]; p < 0.001, Z = −5.93, Wilcoxon sign-rank test). Similar results were obtained from the 14 sessions where topical anesthetic was applied (Fig 3B, green circles): 45 percent of the hippocampus neurons (N = 31/69) were entrained by tACS, and PLVs significantly increased compared to sham (p < 0.001; Wilcoxon sign-rank test; Z = −5.39). Crucially, topical anesthesia did not significantly alter the strength of entrainment during tACS (Fig 3C; p = 0.35; Z = −0.92; Wilcoxon rank-sum test) or the proportion of neurons entrained (p = 0.72, Odds Ratio = 1.19; Fisher’s Exact Test). A block-by-block regression analysis (*Methods* “Topical Anesthesia”) indicates that these results cannot be explained by a decrease in the effectiveness of the anesthesia over time.

**Fig 3.**
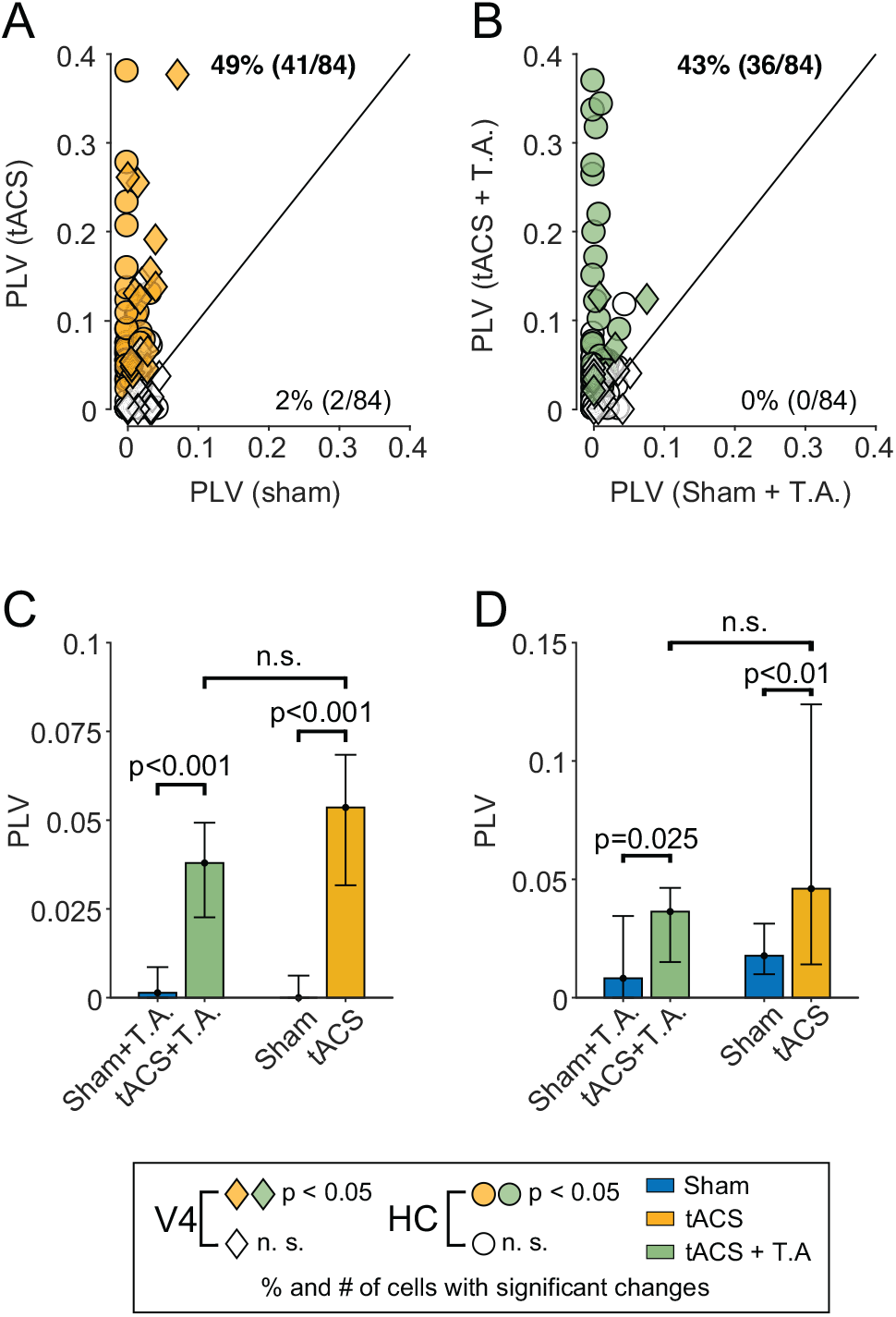
Topical anesthesia does not reduce neuronal entrainment. **(A & B)** PLVs for individual neurons recorded during control (Panel A, yellow) and topical anaesthesia sessions (Panel B, green). Each point compares one neuron’s entrainment during sham stimulation (x-axis) and tACS (y-axis). Circles and diamonds indicate hippocampal and V4 neurons, respectively. Filled shapes indicate neurons that exhibited an individually significant change (p<0.05; randomization test) in entrainment between sham and tACS. PLVs tend to be above the unity line (black), indicating increased entrainment during tACS. The proportion of cells exhibiting significantly increased or decreased entrainment is shown above/below the unity line. Values in bold indicate proportions larger than would be expected by chance (i.e., outside the 95% binomial confidence interval for p=0.05). **(C & D)** Summary plot of the data shown above. Bars indicate median and 95% confidence intervals of median)the data from each condition in the hippocampus (Panel C) and V4 (Panel D); ns: not significant (p>0.05)

Since these negative results may reflect true equivalence between the control and anesthetic conditions or a lack of statistical power, we performed equivalence testing [17]. We found that the difference between the proportion of entrained neurons is significantly equivalent to zero (within ±20%; p = 0.047, Z = −1.67; TOST), suggesting that the results are due to equivalence between the conditions, rather than a lack of statistical power. Similar results were obtained when we lowered the stimulation amplitude to 1 mA, below the animal’s perceptual threshold (Fig 2). As Fig S1 shows, neurons tested during EMLA anesthesia still show significantly increased entrainment during 1 mA tACS (p<0.05; N = 12, randomization tests), and the strength of entrainment was again not significantly different between EMLA and control sessions (p > 0.05; Wilcoxon rank-sum test).

The hippocampus receives input from many brain regions, and may therefore be sensitive to any residual somatosensory input not blocked by the topical anesthesia. We therefore recorded from 43 neurons in area V4, a midlevel visual area on the cortical surface that is less coupled to the somatosensory system [18–20]. For these experiments, the electrode montage was switched to one that optimally stimulated V4, and the current was maintained at ±1 mA to limit the resulting field strength to 1 V/m, which approximates that measured in human cortical areas [21]; this stimulation amplitude also maintains skin stimulation below the animal’s behavioral threshold. As before, tACS increased both the number of neurons entrained and strength of their entrainment. Under control conditions, tACS entrained 46 percent of V4 cells (Fig 3A, diamonds, N = 13/28; p<0.05 per-cell permutation tests) and led to a significant increase in PLV (median sham:0.017, CI:[0.009 – 0.031]; median tACS: 0.046, CI: [0.014 – 0.12]; p < 0.01; Z = −3.10, Wilcoxon sign-rank test). Data collected during topical anesthesia sessions (Fig 3B, diamonds) were similar, with 33% (N = 5/15) of the neurons showing increased entrainment to the tACS and a statistically significant increase in PLVs during tACS as compared to sham stimulation (median sham: 0.008, CI: [0 – 0.035]; median tACS: 0.036 CI: [0.015 – 0.046]; p = 0.025; Z = −2.23, Wilcoxon sign-rank test). Entrainment during EMLA sessions was, again, not significantly different, in terms of median PLV value (p = 0.36; Z = −0.92; Wilcoxon rank-sum) or proportion of neurons entrained (p = 0.52, Odds Ratio = 1.71, Fisher’s Exact Test) between these two conditions (Fig 3D). These results are generally inconsistent with the indirect stimulation hypothesis of tACS.

Another way to test this hypothesis is to examine the stimulation phase at which neurons become entrained. If the electric field were directly causing neuronal entrainment, spikes should be clustered near the peak of the tACS waveform (90°), where the field’s depolarizing effects are strongest. In contrast, indirect peripheral influences would likely arrive at our recording sites at later phases, due to synaptic transmission delays. Our 20 Hz stimulation has a period of 50ms, so only a handful of intervening synapses, each with a few millisecond delay [22, 23], would be sufficient to introduce a lag of 45° (6.25 ms) in the preferred phase. Fig S2 shows the preferred firing phases of the neurons from both areas. During tACS, the preferred firing phases of individual neurons are significantly concentrated around 90° (*V-*test: p = 0.0121 for **Θ_0_**= 90°) during tACS, rather than being shifted towards later phases. Very few neurons had individually significant phase preferences during sham stimulation (Fig 3). However, an analysis of the population phase preference revealed no significant concentration (p = 0.75). Thus, both the phase and strength of entrainment are consistent with direct effects.

## Discussion

These results demonstrate that neuronal entrainment by tACS can survive a topical blockade of somatosensation; indeed for our recording sites, there was little discernible effect of blocking peripheral somatosensory inputs. These data argue strongly against an indirect account of tACS based on entrainment of somatosensory afferents and are more consistent with a direct effect on central neurons.

We note that the field strengths used in these experiments are representative of human tACS. Fields of up to 0.8 V/m have been predicted and measured in human cortex [21]; stronger fields (of up to 2 V/m) may be achievable in other structures, albeit with a multielectrode stimulation montage [24]. With the two-electrode montages used here, fields are strongest near the cortical surface, so other areas may receive stronger stimulation than our hippocampal target. Without recording from a multitude of other cortical areas, it is not possible to rule out indirect entrainment of hippocampal neurons *via the cortex*, but our results suggest that indirect effects from the periphery are unlikely. This is even less likely in V4, a mid-level visual area on the cortical surface that predominately receives visual inputs [18–20].

Previous work has attempted to address the role of somatosensory input by varying the locus of stimulation. We previously reported that entrainment of hippocampal and basal ganglia neurons was abolished by shifting the tACS electrodes to the contralateral hemisphere [3]. Likewise, Johnson, et al. [2] found that shoulder stimulation did not entrain neurons in the pre- and post-central gyrus. However, Asamoah, et al. [5] reported that transcutaneous stimulation of the limbs entrained neurons and EEG in motor cortex. Variations in somatosensory innervation or connectivity between test and control locations have been suggested as a possible explanation for these discrepancies [25], but our experiments allow us to exclude that mechanism by stimulating the same skin locations with and without somatosensory input.

Peripheral nerve stimulation nevertheless has undeniable perceptual consequences that may confound behavioral experiments, and could produce neural effects in areas that receive especially strong somatosensory input. Scalp stimulation may also affect the vagus or other cranial nerves [9], and these mechanisms could even be combined to produce more robust entrainment [26]. These effects will necessarily vary across brain regions and stimulation protocols, frustrating any simplistic attempt at interpretation. Nevertheless, our data, combined with previous work ruling out entrainment via off-target stimulation of the retina [3], support the hypothesis that tACS is capable of directly entraining central neurons.

## Methods

This paper describes data collected from two adult male rhesus monkeys (*Macaca mulatta*), using techniques and experiments that, with the exception of the topical anesthesia, are virtually identical to those in our previous work [3, 16]. Monkey N (7 year old male, 10 kg) participated in the experiments described in Krause, et al. [3] while Monkey S (9 year old male, 20 kg) was obtained specifically for these experiments. All procedures were approved by the Montreal Neurological Institute’s Animal Care Committee (#5031), conformed to the guidelines of the Canadian Council on Animal Care, and were supervised by qualified veterinary staff. When not in the lab, animals were socially housed, received regular environmental enrichment and had access to large play arenas.

### Transcranial Alternating Current Stimulation

Using the method described in our previous work [3, 16, 27], we built an individualized finite-element model from MRIs of the animals’ head and neck, which were solved to find a two-electrode montage that maximized field strength at the recording sites. For the hippocampal recording sites, this montage corresponds to Fp1/O1 in 10-10 coordinates, and was predicted to produce a field of 0.26 V/m when 2 mA of current was applied. We verified this prediction in Krause, et al. [3], and reported that the mean field strength was 0.19 ±0.02 V/m (mean ± standard error) with peak strengths of up to 0.35 V/m. For the V4 site, a montage consisting of Fp1/P7 was found to produce field of ~1 V/m with 1 mA of stimulating current, which approximates field strengths achievable in humans [21].

We applied stimulation using an unmodified StarStim8 system (Neuroelectrics, Barcelona, Spain), using 1 cm (radius) high-definition Ag/AgCl electrodes (PISTIM; Neuroelectrics; Barcelona) coated with conductive gel (SignaGel) and attached to the intact scalp with a thin layer of silicon elastomer (Kwik-Sil, World Precision Instruments). Electrode impedance was continuously monitored during the experiment and was typically between 1-2KΩ, and always below 10 KΩ.

Stimulation consisted of a 20 Hz sinusoidal waveform. We did not optimize the stimulation frequency or phase for individual neurons, but our previous experiments [3] demonstrate that about half of neurons readily entrain to this frequency. For the neurophysiology experiments, current was linearly ramped up from 0 to ±1 or ±2 mA over 10 seconds, held at full amplitude (±1 or ±2 mA) for five minutes, and then ramped back down to 0, again over 10 seconds. The sham stimulation contains the same ramp-up period, but current remained at full amplitude (±1 or ±2 mA) for 10 seconds, before being ramped back down. Since steeper ramps produced stronger percepts, the ramp length was decreased to 2 seconds for the behavioral experiments.

### Topical Anesthesia

Since previous work has used 5% EMLA cream [5, 12, 13, 28] to control somatosensory input during transcranial electrical stimulation, we adopted the same approach. At the beginning of the topical anesthesia experiments, a thick layer of EMLA cream (~3 grams) was applied to a 5 cm (diameter) region surrounding each electrode site. Following the manufacturer’s recommendations, the cream was tightly covered with a plastic dressing and allowed to absorb for approximately one hour (median: 52 minutes, range: 46-72 minutes). The skin was then cleaned with soap and water, followed by alcohol, and allowed to air-dry before the tACS electrodes were attached. Recording typically began 15-20 minutes later.

We used the animal’s behavioral performance to validate the effectiveness of anesthesia. In our previous studies [3, 16], animals were initially distracted and agitated by the onset of tACS, but eventually learned to continue working despite the evoked percepts. If EMLA effectively blocks somatosensation, applying it around tACS electrodes should reduce distractions and increase time spent on-task. Since naive subjects would be most sensitive to these effects, a well-trained monkey that had never received tACS (Monkey S) was used for this experiment. Two pairs of tACS electrodes were placed on its head, one on anesthetized skin over one hemisphere and the other at identical locations on the untreated contralateral side. After a twenty minutes delay that accounted for the setup time needed for neurophysiological experiments, the monkey performed a visual fixation task while bursts of tACS were sporadically applied through each pair of electrodes. As Fig 2 shows, time-on-task was significantly increased at all stimulation amplitudes when tACS was applied to the anesthetized skin, as compared to control sites (0.5mA: p = 0.028; Z = −2.19, 1mA: p = 0.009; Z = −2.60; 2mA: p = 0.016; Z = −2.39; Wilcoxon sign-rank tests). Furthermore, performance was not significantly different from ceiling (100%) during 0.5 and 1.0 mA tACS (0.5 mA: p = 0.07, t(10) = −2.01; 1 mA: p = 0.12; t(10) = −1.72; 1-sample *t*-tests) with anesthesia, but was reduced during control conditions (p = 0.01 t(10) = −3.13 and p<0.001 t(10) = −5.04, respectively). These data suggest that EMLA increases somatosensory thresholds to ~1 mA, and reduces—but does not eliminate—sensation at 2.0 mA, just as it does in humans [5].

Experiments were always completed within the two-hour anesthetic window recommended by the manufacturer (median experiment duration: 68 minutes, range: 24-92 minutes). To verify that the anesthesia remained constant throughout each session, we analyzed each block of data collected during anesthesia sessions separately, using a mixed-effects model with fixed effects of stimulation time (tACS or sham), EMLA ‘dose’ (the duration of EMLA pretreatment, in seconds), and the time elapsed between EMLA removal and midpoint of each block (in seconds). The model also included a random intercept for each block (to account for neurons recorded simultaneously) and a random intercept and slope for stimulation type (to account for cell-specific factors). As in the main text, we find that entrainment significantly increases during tACS (p < 0.001, t(267) = 3.35), but the model’s anesthesia-related coefficients are both indistinguishable from zero (EMLA dose: p = 0.82; t(267) = 0.225; time elapsed: p = 0.49; t(267) = −0.692). These features had little impact on the model, and removing them increased its parsimony, as measured by AIC and BIC (**Δ**AIC = 3.5; **Δ**BIC = 10.7). A similar non-parametric analysis found no significant differences in entrainment between blocks that were recorded 30, 45, or 60+ minutes after EMLA removal (p = 0.97; **Χ^2^**(2) = 0.048; Kruskal–Wallis test).

### Behavioral Task

Since arousal and oculomotor activity can strongly affect neural oscillations, we used a simple fixation task to control the animal’s behavioral state and to minimize eye movements. Animals sat in a standard primate chair (Crist Instruments; Hagerstown, Maryland), 57 centimeters from a computer monitor covering the central 30° × 60° of their visual fields. We monitored eye position non-invasively, using an infrared eyetracker (Eyelink-1000; SR Research, Ontario Canada). Monkeys were trained to fixate a small black target (~0.5°) presented against a neutral grey background. Whenever their gaze remained on this target for ~1-2 seconds, they received a liquid reward. Inter-reward intervals were drawn from an exponential distribution (with a flat hazard function) to prevent entrainment to rewards or expected rewards. Both animals had received extensive training before these experiments, and tended to maintain their gaze continuously on the fixation target. Custom software written in Matlab (The Mathworks, Natick, MA, USA) controlled the behavioral task and coordinated the eye tracker, tES stimulator, and recording hardware.

### Neural Data Collection

Single-neuron data were initially obtained from the left hippocampus, an interesting test-bed for these experiments because it receives input from a wide range of areas, including sensory ones, yet is not tightly tied to any specific modality. Additional data was collected from Area V4, a mid-level visual area that is less coupled to the somatosensory system.

At the start of each recording session, we penetrated the dura with a sharpened 22 ga. stainless steel guide tube. A 32-channel V-Probe with 150 μm spacing (Plexon Inc; Dallas, Texas) was then inserted into the guidetube and positioned with a NaN Microdrive (NaN Instruments; Nazareth Illit, Israel). The target depth was determined from the animals’ MRIs, and, for monkey N, confirmed via CT, as shown in [3].

Wideband signals were recorded using a Neural Interface Processor (Ripple Neuro; Salt Lake City, Utah). Signals were referenced against the guidetube, bandpass filtered between 0.3 and 7500 Hz, and stored at 30,000 Hz with 16 bit/0.21 μV resolution for offline analysis. The raw wideband signals were first preprocessed with a PCA-based filtering algorithm [29] to attenuate stimulation artifacts. Next, single units were identified by bandpass filtering the signal between 500-7000 Hz with a 3^rd^ order Butterworth filter and thresholding it at ±3 standard deviations. The 2 ms segments around each threshold crossing were then sorted using UltraMegaSort 2000, a *k-*means overclustering algorithm [30]. Its results were manually reviewed and refined to ensure that each unit had a clear refractory period, stable width and amplitude, and good separation in PCA space.

### Data Analysis

We quantified the neurons’ entrainment to the electrical stimulation using pairwise phase consistency (PPC), a measure of the synchronization between a point process (spikes) and a continuous signal (tACS or local field potential) with statistical advantages over a direct calculation of phase-locking values or spike-field coherence [31]. The PPC scores are an unbiased estimate of the square of the phase-locking values (PLVs), a more commonly used measure, so we report values as PLVs to facilitate comparison with other work.

Neurons may fire rhythmically even in the absence of stimulation. We therefore compared entrainment to the tACS waveform (during stimulation) with entrainment to the corresponding frequency component (20±1 Hz) of the local field potential (LFP), during sham stimulation. Only the middle four minutes of each block was analyzed, when no current whatsoever was being applied during the sham condition. In both cases, the continuous signal came from an adjacent channel to avoid spectral contamination by the spiking activity [32]. This approach also accounts for physiological distortions of the tACS waveform [33].

Data acquisition or signal processing artifacts could potentially produce the appearance of entrainment. Our previous work [3] describes a number of analyses and controls addressing this issue, using data collected with the same equipment, monkey, and electrodes. We showed that entrainment is unrelated to firing rate or signal amplitude, while neurons’ waveforms remain consistent between tACS and sham conditions, as well as across different phases of the tACS. These analyses suggest that the entrainment seen here is unlikely to be due to technical artifacts.

Our analyses combine all data collected in the same condition, though a block-by-block analysis (see *Topical Anesthesia*, above) yields the same conclusion. Where possible, non-parametric tests were used to avoid distributional assumptions. Randomization tests were used to compare PPC values across tACS and sham conditions; population-level analyses were carried using Wilcoxon rank-sum and sign-rank tests, as appropriate, and 95% confidence intervals for the median were calculated using the formula in [34]. All statistical tests are two-tailed, except where noted. Sample sizes were determined based on our previous work and data was analyzed using Matlab (The Mathworks, Natick, MA) and R [35].

## Acknowledgements

We thank Julie Coursol, Cathy Hunt, and Dr. Fernando Chaurand for outstanding technical assistance and Luiza Volpi for assistance during some recordings. This work was supported by Defense Advanced Research Projects Agency Contract N66001-16-C-4058 (Memory Enhancement by Modulation of Encoding Strength; to C.C.P.), Canadian Institutes of Health Research Grant MOP-115178 (to C.C.P.), and a Jeanne Timmins Costello Postdoctoral Fellowship (to P.G.V). The views, opinions, and/or findings expressed are those of the authors and should not be interpreted as representing the official views or policies of the Department of Defense or the US Government.

## Author Contributions

P.G.V and M.R.K collected and analyzed the data. P.G.V, M.R.K, and C.C.P designed the experiments and wrote the paper.

## Data Availability

The values of individual data points are available in Table S1, S2, and S3. Due to its large size, the raw wideband data is available by request; please contact C.C.P to make arrangements.

## Supporting Information

**Fig S1.**
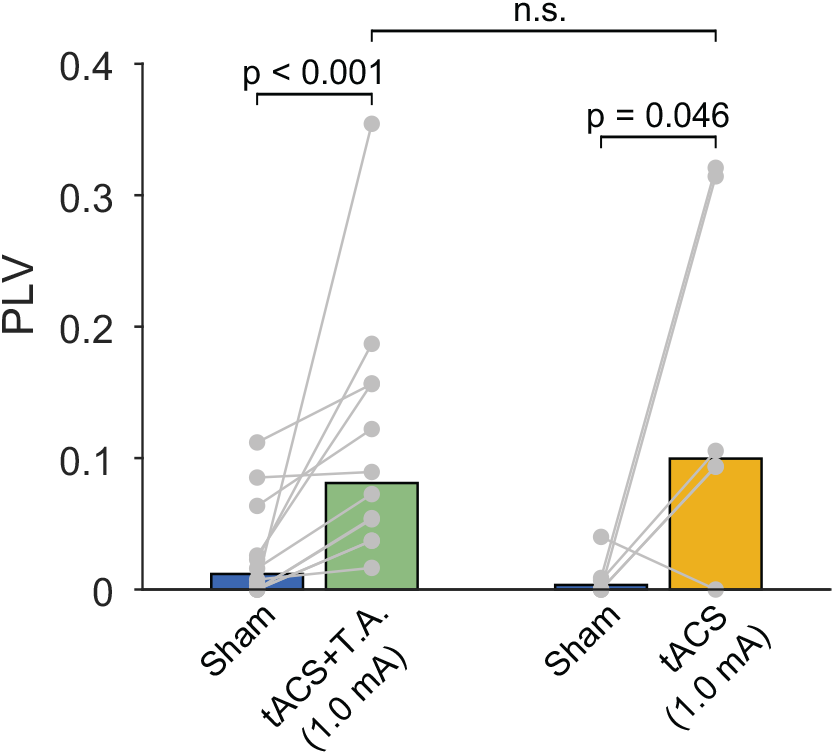
1 mA tACS entrains hippocampal neurons. Data shown in the same style as Fig 3C-D. PLVs for individual neurons are indicated by grey points, with lines connecting observations from the same cell. As before, tACS increases neuronal entrainment during topical anesthesia (TA, green) and control sessions (yellow), compared to the corresponding sham conditions. However, no significant difference (n.s; p>0.05) was detected between tACS and tACS + topical anesthesia.

**Fig S2.**
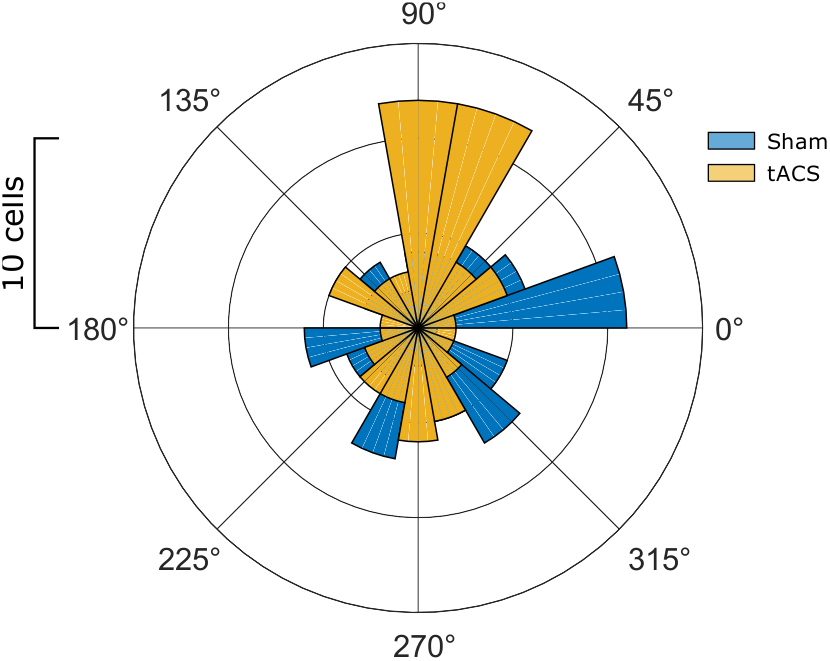
tACS shifts preferred phases to 90°. Distribution of preferred firing phases for the neurons shown in Figure 3 during sham (blue) and 20 Hz tACS (yellow) stimulation. During tACS, neurons preferentially fired near the peak of the tACS waveform (90°) but no such concentration was observed under sham conditions. Note that the preferred phase estimates for sham stimulation are noisy because neurons were minimally entrained to the 20 Hz component of the LFP (Fig 3).

**Table S1. Individual data values for Fig 2.** Each entry indicates the percent of time during which the animal maintained its gaze within 2° of the fixation site. The Data is available in a PDF (Table S1A) and Excel file (Table S1B).

**Table S2. Individual data values for Fig 3 and Fig S2.** Rows in red indicate neurons that showed individually-significant changes in entrainment, shown as filled shapes in Fig 3A and 3B. TA denotes topical anesthesia. Data is available in a PDF (Table S2A) and Excel file (Table S2B).

**Table S3. Individual data values for Fig S1.** Rows in red indicate neurons that showed individually-significant changes in entrainment, shown as filled shapes in Fig 3A and 3B. TA denotes topical anesthesia. Data is available in a PDF (Table S3A) and Excel file (Table S3B).

