## Supplementary material for "tACS entrains neural activity while somatosensory input is blocked": Table S1 (PDF)

| Sample | Time on task (%) | | | | | |
|---|---|---|---|---|---|---|
| | 0.5 mA | | 1 mA | | 2 mA | |
| | tACS | tACS + T.A. | tACS | tACS + T.A. | tACS | tACS + T.A. |
| **1** | 0.00 | 58.22 | 17.71 | 79.44 | 31.14 | 66.29 |
| **2** | 16.89 | 55.92 | 14.97 | 96.88 | 23.35 | 71.54 |
| **3** | 90.10 | 100.00 | 67.14 | 91.37 | 33.87 | 90.61 |
| **4** | 9.97 | 100.00 | 34.27 | 100.00 | 63.94 | 79.93 |
| **5** | 98.22 | 95.40 | 88.22 | 82.81 | 25.41 | 35.77 |
| **6** | 80.31 | 68.20 | 74.65 | 91.86 | 17.29 | 47.08 |
| **7** | 80.33 | 100.00 | 79.71 | 100.00 | 58.76 | 56.30 |
| **8** | 92.53 | 100.00 | 91.07 | 94.33 | 71.10 | 63.39 |
| **9** | 42.47 | 100.00 | 18.07 | 99.26 | 36.56 | 52.70 |
| **10** | 90.93 | 100.00 | 70.71 | 98.77 | 66.25 | 73.49 |
