## Supplementary material for "tACS entrains neural activity while somatosensory input is blocked": Table S2 (PDF)

| Cell | Area | T. A. | tACS current | PLV (Sham) | PLV (tACS) | Phase (Sham) | Phase (tACS) |
|---|---|---|---|---|---|---|---|
| 1 | HC | No | 2 mA | 0.019 | 0.032 | N.A. | N.A. |
| **2** | HC | No | 2 mA | 0.000 | 0.031 | -2.311 | 1.282 |
| 3 | HC | No | 2 mA | 0.034 | 0.000 | N.A. | N.A. |
| 4 | HC | No | 2 mA | 0.000 | 0.008 | N.A. | N.A. |
| **5** | HC | No | 2 mA | 0.013 | 0.104 | 2.573 | -1.133 |
| **6** | HC | No | 2 mA | 0.015 | 0.108 | -3.024 | 0.746 |
| **7** | HC | No | 2 mA | 0.000 | 0.122 | -0.819 | -1.711 |
| 8 | HC | No | 2 mA | 0.034 | 0.070 | N.A. | N.A. |
| 9 | HC | No | 2 mA | 0.013 | 0.015 | N.A. | N.A. |
| 10 | HC | No | 2 mA | 0.000 | 0.000 | N.A. | N.A. |
| **11** | HC | No | 2 mA | 0.000 | 0.044 | 0.238 | 1.600 |
| 12 | HC | No | 2 mA | 0.000 | 0.000 | N.A. | N.A. |
| 13 | HC | No | 2 mA | 0.043 | 0.000 | N.A. | N.A. |
| 14 | HC | No | 2 mA | 0.000 | 0.060 | N.A. | N.A. |
| 15 | HC | No | 2 mA | 0.000 | 0.074 | N.A. | N.A. |
| 16 | HC | No | 2 mA | 0.000 | 0.000 | N.A. | N.A. |
| **17** | HC | No | 2 mA | 0.005 | 0.068 | 1.743 | 1.605 |
| **18** | HC | No | 2 mA | 0.000 | 0.074 | -0.034 | 1.079 |
| **19** | HC | No | 2 mA | 0.020 | 0.086 | -3.033 | 0.592 |
| **20** | HC | No | 2 mA | 0.045 | 0.000 | 0.177 | -2.531 |
| 21 | HC | No | 2 mA | 0.019 | 0.000 | N.A. | N.A. |
| **22** | HC | No | 2 mA | 0.000 | 0.136 | 1.040 | -1.162 |
| 23 | HC | No | 2 mA | 0.013 | 0.027 | N.A. | N.A. |
| **24** | HC | No | 2 mA | 0.000 | 0.206 | 0.147 | -0.545 |
| **25** | HC | No | 2 mA | 0.000 | 0.232 | 0.209 | -0.208 |
| **26** | HC | No | 2 mA | 0.000 | 0.158 | 2.192 | -1.122 |
| **27** | HC | No | 2 mA | 0.000 | 0.088 | 0.010 | 2.314 |
| **28** | HC | No | 2 mA | 0.006 | 0.032 | -2.205 | -0.958 |
| **29** | HC | No | 2 mA | 0.000 | 0.062 | -2.817 | -0.460 |
| **30** | HC | No | 2 mA | 0.000 | 0.380 | 2.582 | 0.162 |
| **31** | HC | No | 2 mA | 0.000 | 0.277 | -1.281 | 1.380 |
| 32 | HC | No | 2 mA | 0.032 | 0.075 | N.A. | N.A. |
| 33 | HC | No | 2 mA | 0.000 | 0.000 | N.A. | N.A. |
| 34 | HC | No | 2 mA | 0.000 | 0.054 | N.A. | N.A. |
| **35** | HC | No | 2 mA | 0.000 | 0.066 | -2.856 | 1.032 |
| **36** | HC | No | 2 mA | 0.033 | 0.130 | -1.922 | -2.595 |
| **37** | HC | No | 2 mA | 0.000 | 0.054 | 1.434 | -3.103 |
| **38** | HC | No | 2 mA | 0.000 | 0.053 | 0.542 | 2.621 |
| 39 | HC | No | 2 mA | 0.000 | 0.000 | N.A. | N.A. |
| 40 | HC | No | 2 mA | 0.000 | 0.033 | N.A. | N.A. |
| **41** | HC | No | 2 mA | 0.000 | 0.068 | -1.862 | 3.065 |
| 42 | HC | No | 2 mA | 0.022 | 0.037 | N.A. | N.A. |
| 43 | HC | No | 2 mA | 0.000 | 0.064 | N.A. | N.A. |
| 44 | HC | No | 2 mA | 0.000 | 0.000 | N.A. | N.A. |
| 45 | HC | No | 2 mA | 0.019 | 0.022 | N.A. | N.A. |
| **46** | HC | No | 2 mA | 0.000 | 0.090 | -2.089 | 0.482 |
| **47** | HC | No | 2 mA | 0.024 | 0.077 | -0.720 | 0.709 |
| 48 | HC | No | 2 mA | 0.000 | 0.000 | N.A. | N.A. |
| **49** | HC | No | 2 mA | 0.000 | 0.107 | -2.586 | 0.612 |
| **50** | HC | No | 2 mA | 0.019 | 0.073 | -2.208 | 0.723 |
| **51** | HC | No | 2 mA | 0.000 | 0.022 | -2.076 | 0.539 |
| 52 | HC | No | 2 mA | 0.010 | 0.000 | N.A. | N.A. |
| **53** | HC | No | 2 mA | 0.000 | 0.048 | -2.667 | 0.302 |
| 54 | HC | No | 2 mA | 0.015 | 0.030 | N.A. | N.A. |
| 55 | HC | No | 2 mA | 0.002 | 0.035 | N.A. | N.A. |
| 56 | HC | No | 2 mA | 0.021 | 0.028 | N.A. | N.A. |
| **57** | V4 | No | 1 mA | 0.010 | 0.131 | -3.066 | -2.934 |
| **58** | V4 | No | 1 mA | 0.007 | 0.041 | -1.132 | 2.470 |
| 59 | V4 | No | 1 mA | 0.034 | 0.000 | N.A. | N.A. |
| 60 | V4 | No | 1 mA | 0.000 | 0.000 | N.A. | N.A. |

| | | | | | | | |
|---|---|---|---|---|---|---|---|
| 61 | V4 | No | 1 mA | 0.035 | 0.014 | N.A. | N.A. |
| 62 | V4 | No | 1 mA | 0.018 | 0.046 | N.A. | N.A. |
| 63 | V4 | No | 1 mA | 0.031 | 0.000 | N.A. | N.A. |
| 64 | V4 | No | 1 mA | 0.017 | 0.017 | N.A. | N.A. |
| **65** | V4 | No | 1 mA | 0.018 | 0.124 | 0.828 | 1.055 |
| **66** | V4 | No | 1 mA | 0.006 | 0.048 | 0.096 | -1.386 |
| **67** | V4 | No | 1 mA | 0.039 | 0.191 | 0.359 | 1.323 |
| 68 | V4 | No | 1 mA | 0.045 | 0.037 | N.A. | N.A. |
| **69** | V4 | No | 1 mA | 0.070 | 0.375 | 0.117 | 1.380 |
| **70** | V4 | No | 1 mA | 0.032 | 0.155 | 0.143 | 1.437 |
| **71** | V4 | No | 1 mA | 0.005 | 0.054 | -2.862 | 1.607 |
| 72 | V4 | No | 1 mA | 0.000 | 0.000 | N.A. | N.A. |
| **73** | V4 | No | 1 mA | 0.028 | 0.046 | -1.839 | 1.252 |
| 74 | V4 | No | 1 mA | 0.013 | 0.022 | N.A. | N.A. |
| 75 | V4 | No | 1 mA | 0.018 | 0.064 | N.A. | N.A. |
| 76 | V4 | No | 1 mA | 0.020 | 0.143 | N.A. | N.A. |
| **77** | V4 | No | 1 mA | 0.014 | 0.255 | -0.850 | 1.369 |
| 78 | V4 | No | 1 mA | 0.000 | 0.000 | N.A. | N.A. |
| **79** | V4 | No | 1 mA | 0.034 | 0.000 | -1.851 | 1.211 |
| 80 | V4 | No | 1 mA | 0.014 | 0.000 | N.A. | N.A. |
| **81** | V4 | No | 1 mA | 0.040 | 0.138 | 1.411 | 1.073 |
| 82 | V4 | No | 1 mA | 0.000 | 0.002 | N.A. | N.A. |
| **83** | V4 | No | 1 mA | 0.028 | 0.065 | 0.209 | 1.500 |
| **84** | V4 | No | 1 mA | 0.000 | 0.261 | -0.616 | 1.709 |
| 85 | HC | Yes | 2 mA | 0.008 | 0.000 | N.A. | N.A. |
| 86 | HC | Yes | 2 mA | 0.035 | 0.000 | N.A. | N.A. |
| **87** | HC | Yes | 2 mA | 0.000 | 0.075 | 2.276 | 1.132 |
| 88 | HC | Yes | 2 mA | 0.020 | 0.002 | N.A. | N.A. |
| **89** | HC | Yes | 2 mA | 0.000 | 0.032 | 1.472 | -1.886 |
| 90 | HC | Yes | 2 mA | 0.000 | 0.000 | N.A. | N.A. |
| 91 | HC | Yes | 2 mA | 0.000 | 0.084 | N.A. | N.A. |
| 92 | HC | Yes | 2 mA | 0.000 | 0.023 | N.A. | N.A. |
| 93 | HC | Yes | 2 mA | 0.000 | 0.000 | N.A. | N.A. |
| 94 | HC | Yes | 2 mA | 0.034 | 0.025 | N.A. | N.A. |
| **95** | HC | Yes | 2 mA | 0.000 | 0.040 | -0.652 | 1.958 |
| 96 | HC | Yes | 2 mA | 0.000 | 0.000 | N.A. | N.A. |
| 97 | HC | Yes | 2 mA | 0.000 | 0.000 | N.A. | N.A. |
| **98** | HC | Yes | 2 mA | 0.014 | 0.025 | -0.974 | 2.481 |
| 99 | HC | Yes | 2 mA | 0.000 | 0.000 | N.A. | N.A. |
| **100** | HC | Yes | 2 mA | 0.011 | 0.031 | -0.478 | 2.469 |
| 101 | HC | Yes | 2 mA | 0.013 | 0.017 | N.A. | N.A. |
| 102 | HC | Yes | 2 mA | 0.035 | 0.046 | N.A. | N.A. |
| 103 | HC | Yes | 2 mA | 0.014 | 0.017 | N.A. | N.A. |
| **104** | HC | Yes | 2 mA | 0.021 | 0.052 | 0.572 | -2.327 |
| **105** | HC | Yes | 2 mA | 0.005 | 0.316 | 1.226 | -1.579 |
| **106** | HC | Yes | 2 mA | 0.000 | 0.336 | 1.223 | 1.727 |
| **107** | HC | Yes | 2 mA | 0.012 | 0.343 | 2.553 | 1.776 |
| **108** | HC | Yes | 2 mA | 0.000 | 0.263 | 2.288 | -1.910 |
| **109** | HC | Yes | 2 mA | 0.000 | 0.369 | -0.951 | -1.482 |
| 110 | HC | Yes | 2 mA | 0.009 | 0.016 | N.A. | N.A. |
| 111 | HC | Yes | 2 mA | 0.000 | 0.012 | N.A. | N.A. |
| **112** | HC | Yes | 2 mA | 0.000 | 0.072 | -1.210 | -0.836 |
| 113 | HC | Yes | 2 mA | 0.008 | 0.025 | N.A. | N.A. |
| **114** | HC | Yes | 2 mA | 0.033 | 0.100 | 0.400 | -2.381 |
| 115 | HC | Yes | 2 mA | 0.028 | 0.048 | N.A. | N.A. |
| 116 | HC | Yes | 2 mA | 0.022 | 0.016 | N.A. | N.A. |
| **117** | HC | Yes | 2 mA | 0.000 | 0.038 | 1.898 | 0.383 |
| 118 | HC | Yes | 2 mA | 0.000 | 0.000 | N.A. | N.A. |
| 119 | HC | Yes | 2 mA | 0.000 | 0.000 | N.A. | N.A. |
| **120** | HC | Yes | 2 mA | 0.022 | 0.053 | -1.170 | 1.697 |
| 121 | HC | Yes | 2 mA | 0.000 | 0.000 | N.A. | N.A. |
| 122 | HC | Yes | 2 mA | 0.000 | 0.015 | N.A. | N.A. |

| | | | | | | | |
|---|---|---|---|---|---|---|---|
| 123 | HC | Yes | 2 mA | 0.000 | 0.000 | N.A. | N.A. |
| **124** | HC | Yes | 2 mA | 0.000 | 0.073 | -0.626 | -1.785 |
| 125 | HC | Yes | 2 mA | 0.000 | 0.013 | N.A. | N.A. |
| 126 | HC | Yes | 2 mA | 0.000 | 0.031 | N.A. | N.A. |
| 127 | HC | Yes | 2 mA | 0.000 | 0.006 | N.A. | N.A. |
| 128 | HC | Yes | 2 mA | 0.000 | 0.067 | N.A. | N.A. |
| 129 | HC | Yes | 2 mA | 0.010 | 0.000 | N.A. | N.A. |
| 130 | HC | Yes | 2 mA | 0.045 | 0.117 | N.A. | N.A. |
| **131** | HC | Yes | 2 mA | 0.004 | 0.120 | -0.406 | 1.577 |
| 132 | HC | Yes | 2 mA | 0.000 | 0.055 | N.A. | N.A. |
| **133** | HC | Yes | 2 mA | 0.016 | 0.054 | -1.641 | -0.948 |
| **134** | HC | Yes | 2 mA | 0.009 | 0.037 | -1.965 | 1.641 |
| 135 | HC | Yes | 2 mA | 0.008 | 0.012 | N.A. | N.A. |
| 136 | HC | Yes | 2 mA | 0.000 | 0.057 | N.A. | N.A. |
| **137** | HC | Yes | 2 mA | 0.014 | 0.039 | 2.401 | 1.094 |
| **138** | HC | Yes | 2 mA | 0.013 | 0.089 | -2.459 | -1.811 |
| 139 | HC | Yes | 2 mA | 0.018 | 0.038 | N.A. | N.A. |
| **140** | HC | Yes | 2 mA | 0.000 | 0.034 | 0.858 | -0.317 |
| **141** | HC | Yes | 2 mA | 0.011 | 0.059 | 1.037 | 2.616 |
| **142** | HC | Yes | 2 mA | 0.000 | 0.049 | 1.086 | 2.322 |
| **143** | HC | Yes | 2 mA | 0.001 | 0.198 | 2.038 | -1.222 |
| **144** | HC | Yes | 2 mA | 0.013 | 0.043 | -2.459 | 2.057 |
| **145** | HC | Yes | 2 mA | 0.020 | 0.042 | 1.062 | -2.308 |
| 146 | HC | Yes | 2 mA | 0.000 | 0.045 | N.A. | N.A. |
| **147** | HC | Yes | 2 mA | 0.004 | 0.170 | 0.471 | -1.611 |
| **148** | HC | Yes | 2 mA | 0.009 | 0.218 | 0.878 | 1.503 |
| 149 | HC | Yes | 2 mA | 0.021 | 0.000 | N.A. | N.A. |
| 150 | HC | Yes | 2 mA | 0.000 | 0.006 | N.A. | N.A. |
| 151 | HC | Yes | 2 mA | 0.003 | 0.001 | N.A. | N.A. |
| **152** | HC | Yes | 2 mA | 0.000 | 0.274 | 0.310 | -1.447 |
| **153** | HC | Yes | 2 mA | 0.000 | 0.150 | 0.253 | 1.607 |
| 154 | V4 | Yes | 1 mA | 0.000 | 0.033 | N.A. | N.A. |
| 155 | V4 | Yes | 1 mA | 0.000 | 0.000 | N.A. | N.A. |
| **156** | V4 | Yes | 1 mA | 0.075 | 0.124 | -1.343 | -2.649 |
| 157 | V4 | Yes | 1 mA | 0.051 | 0.040 | N.A. | N.A. |
| 158 | V4 | Yes | 1 mA | 0.006 | 0.000 | N.A. | N.A. |
| 159 | V4 | Yes | 1 mA | 0.041 | 0.000 | N.A. | N.A. |
| **160** | V4 | Yes | 1 mA | 0.030 | 0.069 | -0.812 | -1.522 |
| **161** | V4 | Yes | 1 mA | 0.000 | 0.034 | -0.837 | -2.265 |
| 162 | V4 | Yes | 1 mA | 0.017 | 0.036 | N.A. | N.A. |
| 163 | V4 | Yes | 1 mA | 0.034 | 0.043 | N.A. | N.A. |
| 164 | V4 | Yes | 1 mA | 0.014 | 0.015 | N.A. | N.A. |
| **165** | V4 | Yes | 1 mA | 0.008 | 0.127 | -1.484 | 2.282 |
| 166 | V4 | Yes | 1 mA | 0.000 | 0.046 | N.A. | N.A. |
| **167** | V4 | Yes | 1 mA | 0.000 | 0.021 | 0.426 | 2.795 |
| 168 | V4 | Yes | 1 mA | 0.000 | 0.039 | N.A. | N.A. |
