## Supplementary material for "tACS entrains neural activity while somatosensory input is blocked": Table S3 (PDF)

| Cell | Area | T. A. | tACS current | PLV (Sham) | PLV (tACS) |
|---|---|---|---|---|---|
| 1 | HC | No | 1 mA | 0.040 | 0.000 |
| **2** | HC | No | 1 mA | 0.009 | 0.321 |
| **3** | HC | No | 1 mA | 0.000 | 0.314 |
| **4** | HC | No | 1 mA | 0.000 | 0.094 |
| **5** | HC | No | 1 mA | 0.000 | 0.093 |
| **6** | HC | No | 1 mA | 0.007 | 0.106 |
| **7** | HC | Yes | 1 mA | 0.008 | 0.016 |
| **8** | HC | Yes | 1 mA | 0.000 | 0.054 |
| **9** | HC | Yes | 1 mA | 0.023 | 0.187 |
| **10** | HC | Yes | 1 mA | 0.004 | 0.354 |
| **11** | HC | Yes | 1 mA | 0.000 | 0.037 |
| **12** | HC | Yes | 1 mA | 0.000 | 0.037 |
| **13** | HC | Yes | 1 mA | 0.112 | 0.156 |
| **14** | HC | Yes | 1 mA | 0.064 | 0.122 |
| **15** | HC | Yes | 1 mA | 0.026 | 0.157 |
| **16** | HC | Yes | 1 mA | 0.016 | 0.073 |
| 17 | HC | Yes | 1 mA | 0.000 | 0.053 |
| 18 | HC | Yes | 1 mA | 0.085 | 0.089 |
